# Testing the effectiveness of a commercially sold probiotic on restoring the gut microbiota of honey bees: a field study

**DOI:** 10.1101/2023.09.13.557574

**Authors:** Megan E. Damico, Burton Beasley, Drew Greenstein, Kasie Raymann

## Abstract

Antibiotic use in apiculture is often necessary to ensure the survival of honey bee colonies. However, beekeepers are faced with the dilemma of needing to combat bacterial brood infections while also knowing that antibiotics kill beneficial bacteria important for bee health. In recent years, bee probiotics have become increasingly purchased by beekeepers because of product claims like being able to “replenish the microbes lost due to agricultural modifications of honey bees’ environment” or “promote optimal gut health.” Unfortunately, these products have little scientific evidence to support their efficacy, and previous lab experiments have refuted some of their claims. Here, we performed hive-level field experiments to test the effectiveness of SuperDFM-HoneyBee™ −the most commonly purchased honey bee probiotic in the United States− on restoring the honey bee gut microbiota after antibiotic treatment. We found slight but significant changes in the microbiota composition of bees following oxytetracycline (TerraPro) treatment and no difference between the microbiota of antibiotic treated bees with or without subsequent probiotic supplementation. Moreover, the microorganisms in the probiotic supplement were never found in the guts of the worker bee samples. These results highlight that more research is needed to test the efficacy and outcomes of currently available commercial honey bee probiotic supplements.

## Introduction

Honey bees are an essential part of the global economy. They play a crucial role in the pollination of crops and contribute over 1.8 billion dollars in revenue to crop production in the United States alone [1]. During critical pollination periods, honey bees are often treated with antibiotics as a preventative measure against bacterial infections. Antibiotics have been used in apiculture for over 60 years in the United States and they are also utilized in apiculture in several other countries. The most commonly applied antibiotic by beekeepers in the United States is oxytetracycline, sold as Terramycin or Terra-Pro, which is primarily used to treat and control *Melissococcus plutonius* and *Paenibacillus larvae*, the causative agents of European and American Foulbrood (EFB and AFB), respectively [2, 3]. Both diseases can be spread from one hive to another when bees drift from an infected hive to an uninfected one, when bees rob an infected hive, or when beekeepers exchange or reuse infected equipment. In large commercial operations where hundreds to thousands of hives are maintained and transported across the United States for pollination services, the threat of disease spread is more severe, which could ultimately result in the loss of hundreds of hives. Additionally, inspecting and diagnosing thousands of hives regularly is almost impossible. Therefore, in the case of large commercial operations, it is common practice to treat (metaphylactically) with antibiotics in the spring and fall and/or before and after entering a highly concentrated pollination site to prevent EFB and AFB diseases.

One major issue with antibiotics is that they target not only pathogens but also beneficial microbes [4]. The honey bee gut microbiome plays an essential role in honey bee health, including immune priming, nutrition and metabolism, growth and development, and protection against pathogens [4–8]. Several studies have shown that exposure to chemicals commonly used in apiculture and agriculture, such as antibiotics and pesticides, disrupt the gut microbiota resulting in decreased survival and increased susceptibility to opportunistic infections [9–17]. Specifically, antibiotic treatment has been linked to weakened honey bee immune responses, indicated by reduced expression of antimicrobial peptides [9]. Reduced antimicrobial peptide expression has been shown to result in increased infection by the fungal parasite Nosema and higher titers of Deformed Wing Virus (DWV) and Israeli Acute Paralysis Virus (IAPV) in honey bees [17]. Antibiotic treatment has also been associated with nutritional deficiencies in honey bees, which are likely caused by perturbation of the gut microbiome [15].

While the use of antibiotics can have negative consequences on bee health, there are currently few alternatives for the treatment and prevention of bacterial infections. In December 2022, an American biotechnology company released a study showcasing the first-ever oral vaccine for *P. larvae* [18]. However, the vaccine is still in the early stages of research and development, has not been thoroughly tested in the field, and only targets the causative agent of AFB. Thus, it is safe to assume that antibiotic use will continue for years to come in commercial beekeeping operations in countries where it is legal. Additionally, the increased use and exposure to agrochemicals (for both native bees and honey bees) is of growing concern and may have indirect impacts on the health and stability of the honey bee gut microbiome [19]. Probiotics, defined as “live microorganisms that, when administered in adequate amounts, confer a health benefit on the host” [20], have risen as a viable, cost-effective option to improve animal health. Although no current evidence shows that probiotics provide health benefits to a healthy host with an undisturbed gut microbiome, probiotic application has been shown to effectively restore microbiota members and microbiome function following antibiotic treatment [21, 22]. Therefore, the use of probiotics could help lessen the adverse effects of antibiotic treatment on honey bees [23]. Currently, there are only a few companies commercially selling probiotics for honey bees (i.e., Fat Bee Probiotics, Durvet, SCD Probiotics, and Strong Microbials), none of which contain native honey bee bacteria. The most popular amongst beekeepers in the United States is Strong Microbials SuperDFM®-HoneyBee™ and it is marketed as a product able to “replenish the microbes lost due to agricultural modifications of honey bees’ environment.” Yet, *all* the products mentioned above are exclusively made of microbes isolated from mammals or the environment that have never been identified in the honey bee gut, and to date, have not been scientifically proven to restore the native honey bee gut microbiome.

Here we tested the effectiveness of SuperDFM®-HoneyBee™ (herein referred to as DFM), the most common marketed honey bee probiotic in the United States, on restoring the honey bee gut microbiota after treatment with Terra-Pro (active ingredient: oxytetracycline). We hypothesized that the gut microbiota of honey bees would not be restored by DFM following exposure to the antibiotics due to the lack of native bee gut or hive-associated microbes present in the probiotic supplement. Although previous lab-based studies have found that oxytetracycline severely disrupts the microbiota of honey bees [11,14], overall, we only observed slight changes in microbiota composition following a single in-hive treatment of TerraPro. This discrepancy is likely due to random sampling of bees with unknown levels of antibiotic exposure in our study. However, as predicted, the microorganisms present in the probiotic supplement were never identified in the guts of any sampled bees, even during active probiotic treatment. These results highlight that more basic research is needed to test the efficacy and outcomes of currently available commercial honey bee probiotic supplements, not only for the sake of honey bees but also beekeepers and the environment.

## Methods

### Sample Collection

Nine hives of *Apis mellifera* (subspecies - Italian honey bees), were created from established hives in early Spring 2019 and kept in an apiary in Dallas, North Carolina. Experimental hives had **not** been treated with antibiotics for at least ten years prior to this experiment, but the hives were treated with the acaricide, Apivar^®^, in Summer 2018, and given an oxalic acid treatment in Winter 2018. No additional treatments for varroa mites or other pests were used during the experiment. Additionally, all new hive materials were used (e.g., boxes, frames, and foundations) to reduce any prior contamination. Hives were also distanced from one another to reduce bees drifting into neighboring hives.

Each hive was comprised of three five-frame deep nucs of approximately equal population size. The nine hives were divided into three experimental groups in June 2019: 1) Control (n=3), 2) Terra-Pro treatment only (n=3), and 3) Terra-Pro treatment followed by DFM probiotic treatment (n=3). The treatment hives (Terra-Pro-Only and Terra-Pro+DFM) were given three applications of Terra-Pro (DC-560, Mann Lake) at 4-5 day intervals following the manufacturer’s instructions. One week after the Terra-Pro application period ended (Week-0), 15 bees per hive were sampled (135 bees total) and group three was given SuperDFM®-HoneyBee™, following the manufacturer’s instructions. On Week-1 and Week-2 after probiotic treatment, 15 bees per hive (control, Terra-Pro, and Terra-Pro+DFM) were sampled per time point (270 bees total). The guts from all sampled bees were extracted immediately upon collection in Dallas, NC using sterile forceps and preserved in 100% ethanol at 4°C until they were transported to and processed in Greensboro, NC. In addition to processing the bee gut samples, DNA was also directly extracted from the SuperDFM-HoneyBee™ powder (in duplicate) using the same methods as above. Although exact hive metrics were not recorded during this study, no overall differences in population size or hive health were noted between treatment and control groups, with the exception of some increased open brood mortality observed in the Terra-Pro and Terra-Pro+DFM hives. However, it must be noted that the increased open brood mortality is a subjective observation that warrants further experimental investigation.

### DNA Extraction and Sequencing

Individual bee gut samples were homogenized and DNA was extracted from the bee guts and DFM probiotic powder using the phenol-chloroform with bead beating extraction protocol described in [24]. For the extracted DNA, a two-step 16S rRNA library preparation was performed. The first step consisted of PCR amplification of the V4 region of the 16S rRNA gene using the 515F and 806R primers containing Illumina platform-specific sequence adaptors: Hyb515F_rRNA: 5′-TCGTCGGCAGCGTCAGATGTGTATAAGAGACAGGTGYCAGCMGCCGCGGTA -3′ and Hyb806R_rRNA: 5′-GTCTCGTGGGCTCGGAGATGTGTATAAGAGACAGGGACTACHVGGGTWTCTAAT-3′. The PCR cycling conditions were as follows, 98°C for 30s followed by 20 cycles of 98°C (10s), 58°C (30s), 72°C (30s), with a final extension at 72°C for 7m and a hold at 4°C. The PCR product was cleaned using the AxygenTM AxyPrep Mag™ PCR Clean-up Kit. Samples were indexed using the Illumina Nextera XT Index kit v2 sets A and D. The PCR cycling conditions were 98°C for 2m followed by 15 cycles of 98°C (10s), 55°C (30s), 72°C (30s), with a final extension at 72°C for 7m and a hold at 4°C. The indexed product was cleaned using the AxygenTM AxyPrep Mag™ PCR Clean-up Kit, quantified with a Qubit3.0 (Life Technologies) with the Qubit dsDNA BR Assay kit, and pooled in equal concentrations for sequencing. A 30% PhiX spike-in was included in the pooling library before sequencing to increase the diversity of the run. The amplicon sequencing was completed at UNC Greensboro using an Illumina iSeq100 with 2×150 paired-end reads. Of the 405 bees collected in this study, 376 were successfully sequenced and used for downstream analyses.

### Sequence Analysis

The forward and reverse reads were merged using FLASH v1.2.10 [25] with a minimum overlap of 5 bp, which resulted in a total of 12,242,419 total reads. Joined reads were quality filtered in Qiime2 v 2023.7 [26] using the DADA2 [27] pipeline, resulting in 7,525,882 reads. The data was filtered to remove all sequences corresponding to mitochondria, chloroplast, and unassigned taxa as well as reads present at less than 0.1% frequency. After quality-filtering a total of 6,599,366 reads were retained, with a mean frequency of 12,490 per sample corresponding to 1,298 amplicon sequence variants (ASVs). Our negative control only contained 92 reads and was removed from further analysis. Downstream analyses were performed in Qiime2 v 2023.7 at a sampling depth of 3,000 reads per sample. This sampling depth was chosen to maximize the number of samples included in the analysis while still maintaining enough reads per sample to capture the richness of the dataset. Rarefying to 3,000 reads per sample resulted in retaining 351 samples, including the two DFM powder controls.

### Quantitative PCR

We amplified total copies of the 16S rRNA using the primers EUB338F and EUB518R (Table 1) on an Applied Biosystems QuantStudio 6 Real-Time PCR system. Reactions were completed in triplicate using 5 μL of universal SYBR Green (Bio-Rad, Inc.), 2 μL of molecular grade water, 1 μL (each) of 3 nM primers, and 1 μL of template DNA. The PCR cycle was 95°C (3 min) followed by 40 cycles of 95°C (3 s) and 60 °C (20 s). The average Ct (cycle threshold) was determined for each sample. A gBlocks™ gene fragment (Integrated DNA Technologies) was used to create a standard curve (Table 1) of known copy number. A gBlock™ was used as it allows for more accurate and simple copy number calculations [28]. We then estimated absolute copy number by interpolating the Ct value into standard curves of known copy numbers, from 10^2^–10^8^ copies.

**Table 1.**
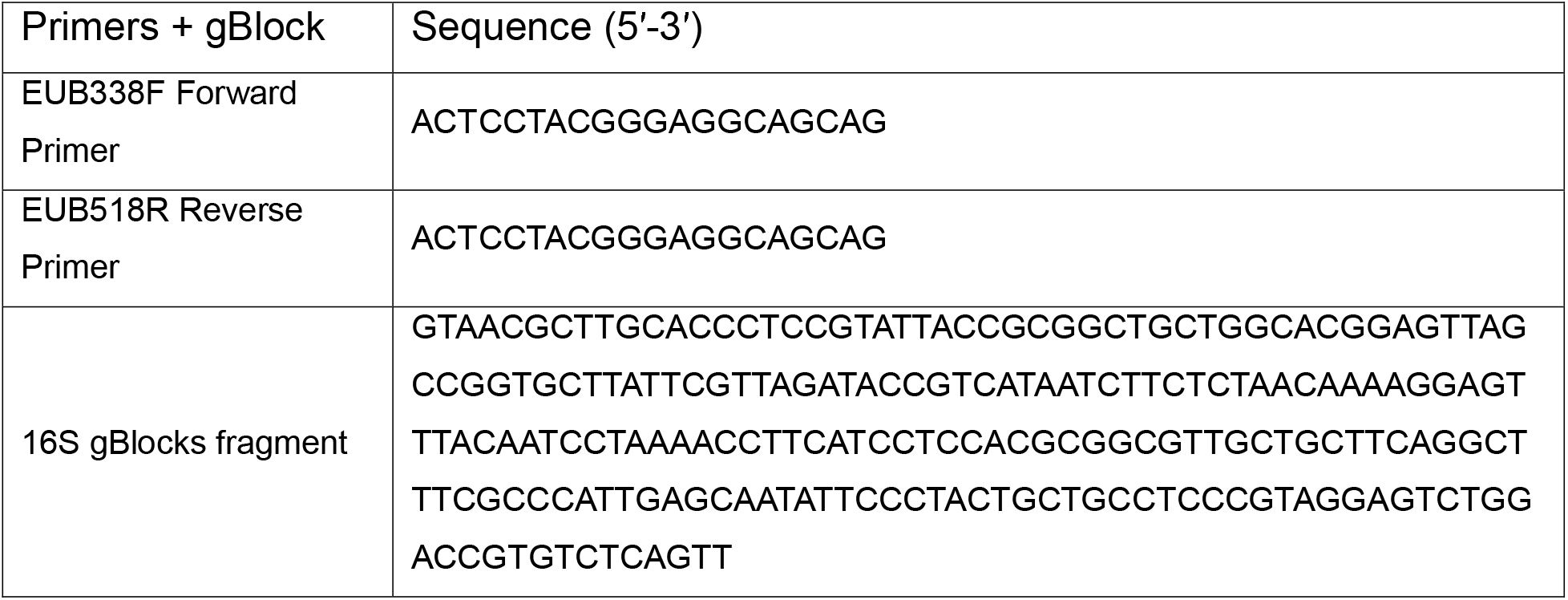
Primers and gBlocks fragment used for qPCR.

### Statistical analysis and data visualization

The script “qiime diversity core-metrics-phylogenetic” was used to perform Alpha and Beta diversity analyses [26]. The taxonomy of the representative sequences was determined using “qiime feature-classifier classify-sklearn” [24] using a classifier trained on the SILVA 16S v138.1 [29] reference database and the BEExact v2023.01.30 [30] reference database. Taxonomic assignments for poorly classified ASVs were confirmed through manual verification using nucleotide BLAST [31] on the NCBI server [32]. The analysis of taxonomic diversity was carried out at both the genus and ASV level. Alpha diversity was statistically tested using the Kruskal-Wallis test with Benjamini-Hochberg FDR correction, and the results were plotted using GraphPad Prism v9.5.1. Beta diversity analyses were done using weighted UniFrac [33, 34] and statistically analyzed using PERMANOVA (999 permutations) with Benjamini-Hochberg FDR correction. The PCoA plots with 95% confidence intervals (stat_ellipse) were generated using Qiime2R [35]. Differences in absolute abundance based on qPCR were determined using the Kruskal-Wallis test with Benjamini-Hochberg FDR correction, and the graph was made in GraphPad Prism v9.5.1.

## Results

In this study, nine honey bee hives (three-story, five-frame deep nucs) were split into three groups (Control, Terra-Pro, and Terra-Pro+DFM). Using 16S rRNA sequencing, we profiled the guts of worker honey bees at each sampling point within the three groups (Figure 1). Finally, we used qPCR to measure the absolute abundance of bacteria to determine if the overall bacterial load differed following antibiotic treatment o after the application of the probiotic supplement.

**Figure 1.**
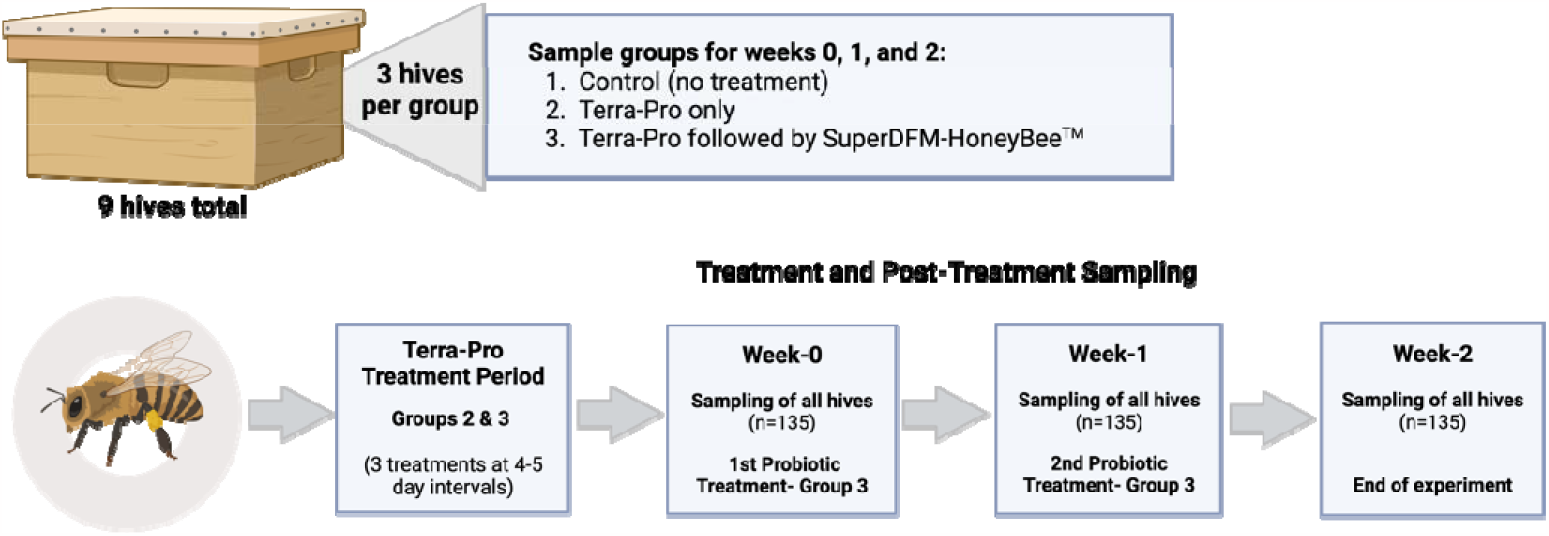
Schematic of experimental setup and sampling timeline. Created with BioRender.com.

### Microbiota diversity analyses of bees from Post-Terra-Pro Pre-DFM treated hives

The first sampling occurred after three antibiotic treatments at 4-5 day intervals had been administered to the Terra-Pro and Terra-Pro+DFM hives (Figure 1; Week-0). At this timepoint the Terra-Pro and Terra-Pro+DFM hives were biological replicates, so we expected them to differ from the Control hives but not from each other. When assessing Alpha diversity, we observed no significant differences in microbiota richness (ASVs) between any of the treatment groups (Figure 2A). Significant differences in microbiota evenness were found when comparing bees from Terra-Pro and Terra-Pro+DFM hives (*Q=*0.01), but not bees from Control and Terra-Pro+DFM hives (*Q=*0.41) or Terra-Pro and Terra-Pro+DFM hives (Q=0.07; Figure 2A). Phylogenetic diversity in the microbiota was significantly higher (even when extreme outliers were excluded) in bees from Control hives than in Terra-Pro or Terra-Pro+DFM hives (*Q=*0.02 and *Q=*0.05, respectively). Phylogenetic diversity did not differ between bees from Terra-Pro and Terra-Pro DFM hives (*Q=*0.76; Figure 2A). Despite minimal differences in Alpha diversity, Beta diversity analysis using weighted UniFrac [34] to analyze microbiota community similarity, revealed that the microbiota composition of Control bees differed significantly from Terra-Pro and Terra-Pro+DFM bees (*Q=*04), whereas bees from the Terra-Pro and Terra-Pro+DFM hives did not significantly differ from each other (*Q=*0.57; Figure 2A). Results indicate that the microbiota was slightly perturbed in the antibiotic treated hives at the end of a standard treatment with Terra-Pro.

**Figure 2.**
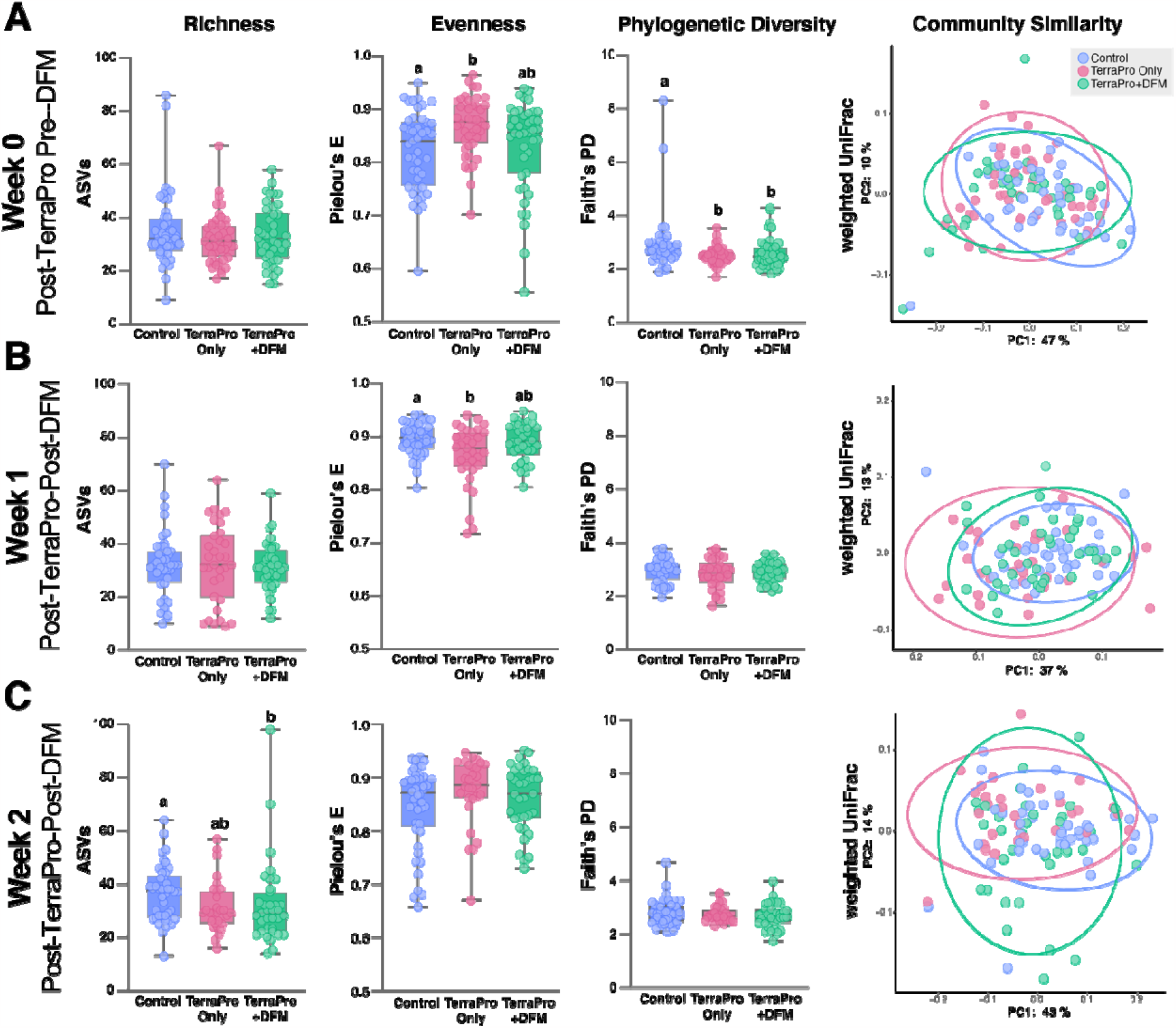
Alpha and Beta diversity comparisons of the gut microbiota of control, Terra-Pro-Only, and Terra-Pro+DFM bees at timepoints A) Week-0 post-Terra-Pro pre-DFM, B) Week-1 post-Terra-Pro post-DFM. And c) Week-2 post-Terra-Pro post-DFM. Alpha diversity metrics were based on richness (# of ASVs), evenness (Pielou’s Evenness index), and phylogenetic diversity (Faith’s Phylogenetic Diversity index). Beta diversity was based on community similarity (weighted UniFrac) and visualized via principal coordinate analysis (PCoA) plots. Alpha diversity significance (*P-value*) was determined using the Kruskall-Wallis test with Benjamini-Hochberg FDR correction (*Q-value*). Significance (*Q*<0.05) is represented by differing letters (absence of letters indicates no significance). For Beta diversity comparisons significance was tested using PERMANOVA with 999 permutations followed by Benjamini-Hochberg FDR correction. Ellipses represent the 95% confidence interval.

### Microbiota diversity analyses of bees from Post-Terra-Pro Post-DFM treated hives

After bees were sampled from each experimental hive post-Terra-Pro treatment (Week-0), the Terra-Pro+DFM hives were immediately administered a treatment of DFM. One week following DFM application to the Terra-Pro+DFM hives, all hives were sampled (Figure 1; Week-1). On Week-1, Alpha and Beta diversity measurements were virtually identical to the post-Terra-Pro treatment but Pre-DFM treatment (Week-0) results. However, no significant differences were found in the evenness of the microbiota across experimental groups (Figure 2B). Additionally, the microbiota composition (based on weighted UniFrac) of Terra-Pro and Terra-Pro+DFM bees differed more from control bees at week-1 (*Q=*0.01) than they did at week-0 (*Q=*0.04). Again, no significant difference was found between the microbiota composition of Terra-Pro and Terra-Pro+DFM bees (*Q=*0.17; Figure 2B). Findings suggest that a single probiotic treatment with DFM did not significantly impact the recovery of the microbiota honey bees treated with Terra-Pro.

Following Week-1 sampling, the Terra-Pro+DFM hives were given a second treatment of DFM, and all hives were subsequently sampled one week later (Figure 1; Week-2). One week after the *second* application of DFM in the Terra-Pro+DFM hives, bees from the Terra-Pro+DFM hives showed a significant decrease in the number of ASVs when compared to Control bees (*Q=*0.05; Figure 2C)). The number of ASVs did not significantly differ between Control and Terra-Pro (*Q=*0.06) or Terra-Pro and Terra-Pro+DFM bees (*Q=*0.72; Figure 2C). No differences in microbiota evenness or phylogenetic diversity were observed between any of the experimental groups at Week-2 (Figure 2C). The microbiota composition (based on weighted UniFrac) of Terra-Pro and Terra-Pro+DFM bees still did not differ (*Q*=0.12), but they remained significantly different from the Control bees at Week-2 (*Q=*0.04). Results indicate that a second probiotic treatment with DFM did not significantly impact the recovery of the microbiota following Terra-Pro treatment.

To access variation across hives *within* experimental groups, we compared the Alpha diversity (i.e., microbiota richness, phylogenetic relatedness, and evenness) of the microbiota of bees from different hives within the same experimental group. Overall, very little variation in Alpha diversity occurred across hives *within* experimental groups at the same sampling timepoint (Figure S1; Dataset S1). We also compared microbiota similarity across hives *within* experimental groups and found that while significant variation occurred across sampling timepoints, variation in Beta diversity between hives at the same timepoint was only observed for Terra-Pro Week-2 Hive-2 and Hive-3 (Figure S2; Dataset S1).

We quantified the absolute abundance of bacteria based on 16S rRNA gene copy number within individual bees from each experimental group at Weeks -0, -1, and -2. At Week-0 (Post-Terra-Pro but pre-DFM treatment), bacterial abundance did not significantly differ between any of the experimental groups (Figure 3). Following the first DFM treatment (Week-1), bees from the TerraPro hives had higher bacterial abundance in their guts than the Control hives (*Q=*0.002). Bacterial abundance did not differ between TerraPro+DFM and Control bees (*Q=*0.36) or the Terra-Pro and TerraPro+DFM bees (*Q=*0.22) at Week-1 (Figure 3). On Week-2, one week following the second application of DFM, bacterial abundance was lower in bees from the Terra-Pro+DFM hives than in bees from the Control (*Q=*0.005) and Terra-Pro (*Q=*0.0002) hives (Figure 3). However, the Terra-Pro and Control bees did not significantly differ in bacterial abundance (*Q=*0.64) at Week-2. These findings suggest that DFM treatment could lead to a reduced number of gut microbes in bees when administered after Terra-Pro.

**Figure 3.**
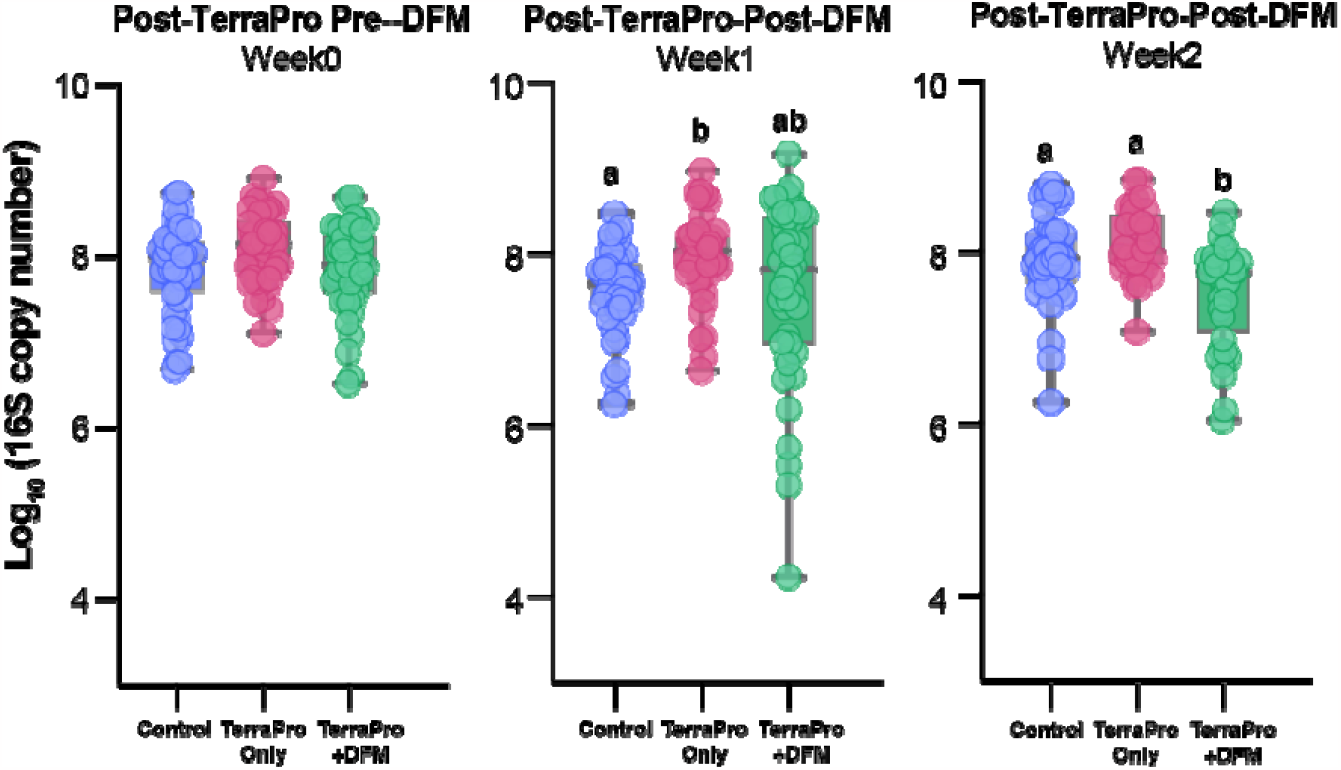
Total bacterial 16S rRNA gene (log_10_ gene copies) copy number estimated by qPCR in Control, Terra-Pro, and Terra-Pro+DFM bees at Week-0, -1, and -2. Box plots show median values with standard deviation and each point represents an individual bee. Significance (*P-value*) was tested using the Kruskal Wallis Test with Benjamini-Hochberg FDR correction (*Q-value*) and is indicated by differing letters.

To visualize the taxonomic changes in microbiota composition across experimental groups and sampling weeks, we averaged the relative abundance of taxa present in all bees from each timepoint and experimental group (Figure 4; see Figure S3 and Dataset S2 for individual bee data). Regardless of treatment group, all honey bees contained the five core honey bee gut microbiome taxonomic groups, *Lactobacillus* Firm4 and Firm5, *Bifidobacteria, Gilliamella*, and *Snodgrassella* (Figure 4). However, shifts in relative abundance were seen amongst all five core taxa and in other less abundant native bee gut members like *Frischella* and *Bartonella within* and *across* all experimental groups at all timepoints (Figure 4). Some notable differences in relative abundance between the controls and treatment groups were an increase in “other” taxa in both the Terra-Pro-Only and Terra-Pro+DFM at Week-1, a decrease in *Snodgrassella* in the Terra-Pro and Terra-Pro+DFM bees at Week-2, and an increase in *Lactobacillus* in the Terra-Pro bees at Week-2 (Figure 4). It is important to note that the SuperDFM-Honeybee™ probiotic bacteria were never found in any of the bee gut samples, even during active treatment with the probiotic powder (Figure 4, Figure S3).

**Figure 4.**
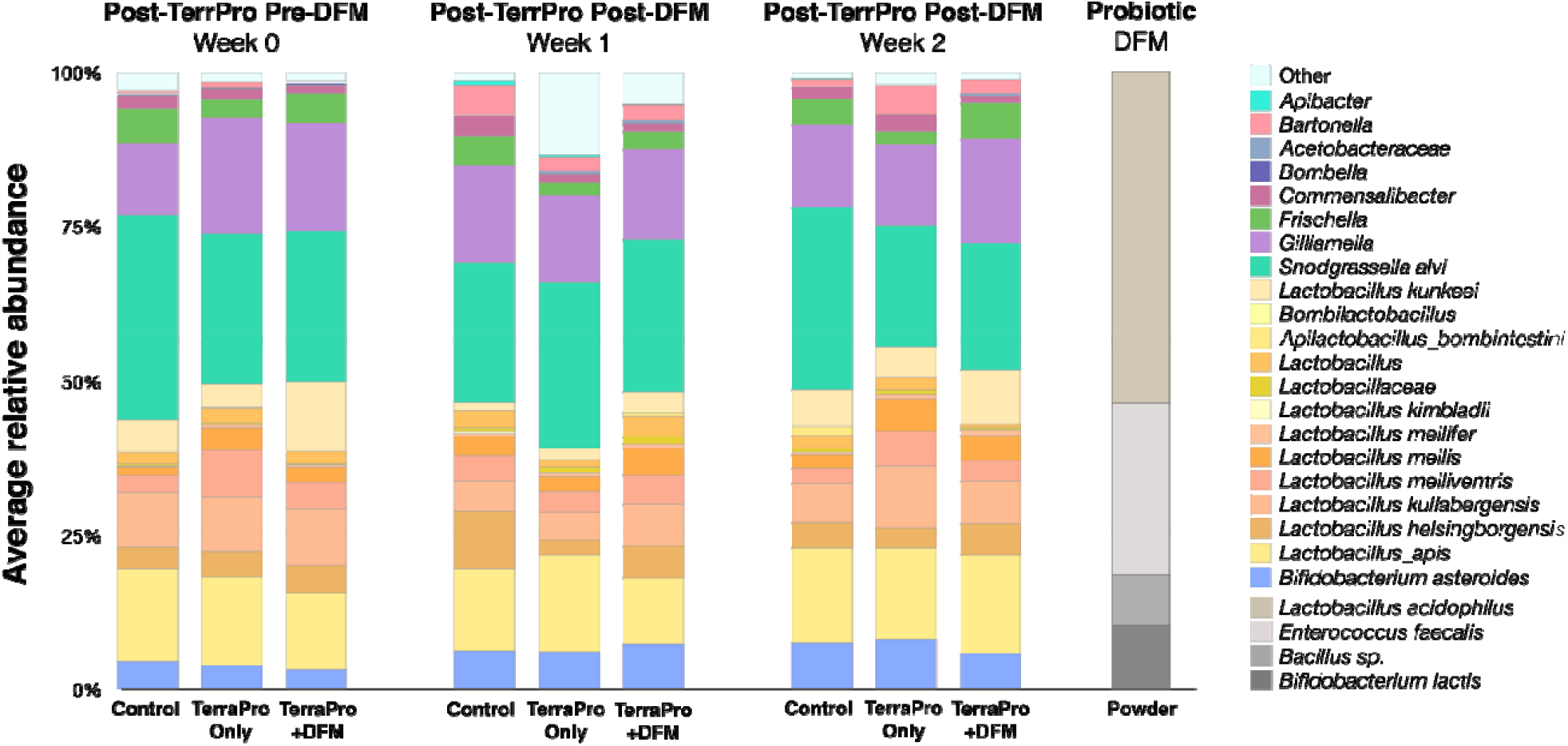
Bar plots representing the average relative abundance of taxa found in the bee gut samples within each experimental group at each sampling timepoint. The DFM powder (probiotic) was also sequenced as a positive control. See Figure S3 for data on individual bees.

## Discussion

Despite promising claims, there is no scientific evidence demonstrating the effectiveness of current commercially available probiotics in restoring the honey bee gut microbiome after a perturbation. Additionally, none of the current commercially available probiotics contain native bee microbes. One of the most popular honey bee probiotics is SuperDFM-HoneyBee™, made by the company Strong Microbials. This product is advertised as a probiotic that can “replenish the microbes lost due to agricultural modifications of honey bees’ environment” and “Promotes optimal gut health”.

Therefore, we sought to determine whether this probiotic can restore the honey bee gut microbiota after a routine treatment with the antibiotic oxytetracycline (Terra-Pro). We found that bees from hives given the probiotic following Terra-Pro treatment were not any more similar to bees from the Control hives than bees from hives only given Terra-Pro treatment. In general, both the Terra-Pro-Only and Terra-Pro+DFM bees displayed more variation between individuals than the Control bees at all sampling timepoints. Moreover, we never detected the microbes present in the probiotic in the guts of bees during or following supplementation and our results suggest that DFM treatment could potentially lead to decreased bacterial abundance (TerraPro+DFM bees had fewer bacteria in their guts after two DFM treatments than Control and Terra-Pro bees). These results are consistent with a previous study that evaluated an unnamed commercial probiotic and revealed that it never colonized the honey bee gut (even of microbiota-depleted bees) and that the probiotic led to an increase in the abundance of opportunistic pathogens [23]. However, given that we only observed a mild amount of microbiota disturbance following TerraPro treatment, we argue that additional hive-level studies are needed before definitive conclusions can be made about the potential health benefits and safety of currently available probiotics.

Although we observed significant differences between control hives and hives treated with Terra-Pro, the effects of the antibiotic treatment were much less severe than expected. This might sound like a promising result for bees and beekeepers, but we are skeptical our study accurately reflects the impacts of antibiotic treatment on the honey bee microbiome, especially because these results contradict several published studies. For example, previous hybrid lab- and field-based studies found that both oxytetracycline (the active ingredient in Terra-Pro) and tylosin (the active ingredient in Tylan) severely perturbed the gut microbiome of bees by reducing the size and composition of bacteria present in the community, resulted in increased susceptibility to opportunistic pathogens, and reduced survivorship [11, 16]. We hypothesize that several factors limited our ability to capture the full effects of Terra-Pro on the honey bee microbiome in our study: 1) bees were only sampled from brood frames at random so we are unsure how long they were exposed to the antibiotic and some could have been newly emerged (lacking the characteristic microbiome), 2) all hives were treated with the acaricide Apivar (active ingredient amitraz) prior to the experiment, which could have impacted their microbiomes [36], and 3) our limited sample size per hive might not have been representative of the hive populations. Therefore, we argue that more extensive and controlled hive-level studies are required to make a definitive statement about the effects of in-hive Terra-Pro treatment as well as the DFM probiotic on the microbiome and health of honey bees.

Recently, several research groups have been investigating other probiotic treatments, which are not yet commercially available to beekeepers. A honeybee-specific lactic acid bacteria (hbs-LAB) probiotic has been developed which contains a mixture of crop and gut associated *Lactobacillus* and *Bifodobacterium* species isolated from honey bees [37, 38]. The hbs-LAB (also referred to as SymBeeotic), has been shown to inhibit *P. larvae* infection when tested in the lab [39, 40] but not at the colony level [41]. To our knowledge SymBeeotic has not been tested for its ability to restore the microbiota after perturbation. Another honey bee probiotic that is currently being tested is the BioPatty™ (also referred to as LX3), which is made of a mixture of three *Lactobacillus* strains, including *Lactobacillus* (*Apilactobacillus) kunkeei* (a hive-associated bacterium that is often found in the honey bee gut) [42–44]. The BioPatty™ has been shown to increase the expression of bee immunity genes, reduce infection of *P. larvae* in infected colonies, and help restore the microbiome following antibiotic perturbation [42–44]. Another research group has been exploring the use of a cocktail of core native bee gut strains as a potential probiotic therapy [16, 23]. This cocktail has been demonstrated to successfully colonize and persist in germ-free bees and reduce the ability for opportunist bacteria to proliferate [16, 23].

Given that all honey bees possess a highly conserved microbiome composed of five core bacterial taxa that are functionally important for be health [4], a probiotic comprised of native honey bee bacteria would increase the chance restoration of the community and eliminate the risk of introducing foreign microbes into the hive ecosystem [23] . However, an additional and important factor to consider when developing honey bee probiotics is how the probiotic should be delivered. In fact, very few studies have been performed to investigate how the mode of delivery of probiotics could impact their effectiveness. One recent study evaluated the impact of delivery method by comparing an LAB (LX3) infused pollen patty vs. a spray-based formulation [44]. Results showed both delivery methods facilitate viable uptake of the LX3 probiotic in adult honey bees, although the strains do not colonize long-term [44]. With this in mind, the continued development and testing of probiotic blends containing native honey bee gut bacteria and an optimal method of administration will hopefully lead to an effective and safe commercially available supplement that can help improve bee and hive health.

## Conclusion

Probiotics, in theory and concept, are a promising solution to enhance bee health, but the current market available products for beekeepers are making claims that far outreach the ability of their products [45]. However, research into proprietary blends of native bee gut bacteria shows promise and holds great potential for revitalizing honey bee health. Along with scientific research, there is a need for policies and regulations that better reflect the growing research on the potentially hazardous impacts of probiotics on honey bee health. Currently, probiotics are regulated as food, which circumvents the responsibility of companies to provide data to the FDA on the product’s long-term safety for the targeted organism and claims made about the product’s effects [46, 47]. The FDA’s Center for Veterinary Medicine (CVM) should revise its policy on Animal Foods with Drug Claims to regulate honey bee probiotics as a nutritional ingredient (which is regulated as a drug) rather than an animal feed (which is regulated as food) [47]. This policy change would require more product testing before commercialization and would save farmers and beekeepers from purchasing ineffective products and help safeguard honey bee health.

## Acknowledgements

This work was supported by the National Science Foundation under grant DEB-1930776 awarded to K.R. We would also like to thank Drs. Louis-Marie Bobay, Heather Hopkins, and John A. Roque III for their constructive feedback and editing assistance on the manuscript.

## Author contributions

M. E. D. performed molecular experiments, collected molecular data, analyzed the data, and wrote and edited the manuscript. D. G. performed molecular experiments and helped edit the manuscript. B. B. helped conceive and organize the project, established honey bee hives, performed all fieldwork, and collected samples. K.R. helped conceive and organize the project, funded the project, analyzed data, created the figures, and wrote and edited the manuscript. All authors contributed critically to the drafts and gave final approval for publication.

## Data availability statement

All raw sequencing reads are deposited in NCBI Sequence Read Bioproject under accession number PRJNA971198. All other data generated are included in the supplemental material files.

## References

1. Keel CC, United States Department of Agriculture (2022) The buzz about pollinators. https://www.usda.gov/media/blog/2022/06/22/buzz-about-pollinators.

2. Mueller CM, Ashley N. Mortensen Cameron J. Jack, Jamie Ellis. Identification and treatment of European Foulbrood in honey bee colonies. https://edis.ifas.ufl.edu/publication/IN1272.

3. Genersch E (2010) American Foulbrood in honeybees and its causative agent, Paenibacillus larvae. J Invertebr Pathol 103:S10–S19. 10.1016/j.jip.2009.06.015

4. Raymann K, Moran NA (2018) The role of the gut microbiome in health and disease of adult honey bee workers. Curr Opin Insect Sci 26:97–104. 10.1016/j.cois.2018.02.012

5. Bonilla-Rosso G, Engel P (2018) Functional roles and metabolic niches in the honey bee gut microbiota. Curr Opin Microbiol 43:69–76. 10.1016/j.mib.2017.12.009

6. Kwong WK, Mancenido AL, Moran NA (2017) Immune system stimulation by the native gut microbiota of honey bees. R Soc Open Sci 4:170003. 10.1098/rsos.170003

7. Zheng H, Nishida A, Kwong WK, et al (2016) Metabolism of toxic sugars by strains of the bee gut symbiont Gilliamella apicola. mBio 7:e01326–16 10.1128/mBio.01326-16

8. Zheng H, Powell JE, Steele MI, et al (2017) Honeybee gut microbiota promotes host weight gain via bacterial metabolism and hormonal signaling. Proc Natl Acad Sci USA 114:4775–4780. 10.1073/pnas.1701819114

9. Kakumanu ML, Reeves AM, Anderson TD, et al (2016) Honey bee gut microbiome Is altered by in-hive pesticide exposures. Front Microbiol 7:1225. 10.3389/fmicb.2016.01255

10. Motta EVS, Raymann K, Moran NA (2018) Glyphosate perturbs the gut microbiota of honey bees. Proc Natl Acad Sci USA 115:10305–10310. 10.1073/pnas.1803880115

11. Raymann K, Shaffer Z, Moran NA (2017) Antibiotic exposure perturbs the gut microbiota and elevates mortality in honeybees. PLoS Biol 15:e2001861. 10.1371/journal.pbio.2001861

12. Motta EVS, Moran NA (2020) Impact of glyphosate on the honey bee gut microbiota: effects of intensity, duration, and timing of exposure. mSystems 5:e00268–20. 10.1128/mSystems.00268-20

13. Motta EVS, Mak M, De Jong TK, et al (2020) Oral or topical exposure to glyphosate in herbicide formulation impacts the gut microbiota and survival rates of honey bees. Appl Environ Microbiol 86:e01150–20. 10.1128/AEM.01150-20

14. Raymann K, Bobay L, Moran NA (2018) Antibiotics reduce genetic diversity of core species in the honeybee gut microbiome. Mol Ecol 27:2057–2066. 10.1111/mec.14434

15. Duan X, Zhao B, Jin X, et al (2021) Antibiotic treatment decrease the fitness of honeybee (Apis mellifera) larvae. Insects 12:301. 10.3390/insects12040301

16. Powell JE, Carver Z, Leonard SP, Moran NA (2021) Field-realistic tylosin exposure impacts honey bee microbiota and pathogen susceptibility, which is ameliorated by native gut probiotics. Microbiol Spectr 9:e0010321. 10.1128/Spectrum.00103-21

17. Deng Y, Yang S, Zhao H, et al (2022) Antibiotics-induced changes in intestinal bacteria result in the sensitivity of honey bee to virus. Environ pollut 314:120278. 10.1016/j.envpol.2022.120278

18. Dickel F, Bos NMP, Hughes H, et al (2022) The oral vaccination with Paenibacillus larvae bacterin can decrease susceptibility to American Foulbrood infection in honey bees—A safety and efficacy study. Front Vet Sci 9:946237. 10.3389/fvets.2022.946237

19. Daisley BA, Chmiel JA, Pitek AP, et al (2020) Missing microbes in bees: how systematic depletion of key symbionts erodes immunity. Trends Microbiol 28:1010–1021. 10.1016/j.tim.2020.06.006

20. Hill C, Guarner F, Reid G, et al (2014) The International Scientific Association for Probiotics and Prebiotics consensus statement on the scope and appropriate use of the term probiotic. Nat Rev Gastroenterol Hepatol 11:506–514. 10.1038/nrgastro.2014.66

21. Beck LC, Masi AC, Young GR, et al (2022) Strain-specific impacts of probiotics are a significant driver of gut microbiome development in very preterm infants. Nat Microbiol 7:1525–1535. 10.1038/s41564-022-01213-w

22. Korpela K, Salonen A, Vepsäläinen O, et al (2018) Probiotic supplementation restores normal microbiota composition and function in antibiotic-treated and in caesarean-born infants. Microbiome 6:182. 10.1186/s40168-018-0567-4

23. Motta EVS, Powell JE, Leonard SP, Moran NA (2022) Prospects for probiotics in social bees. Philos Trans R Soc Lond B Biol Sci 377:20210156. 10.1098/rstb.2021.0156

24. Evans JD, Schwarz RS, Chen YP, et al (2013) Standard methods for molecular research in Apis mellifera. Journal of Apicultural Research 52:1–54. 10.3896/IBRA.1.52.4.11

25. Magoč T, Salzberg SL (2011) FLASH: fast length adjustment of short reads to improve genome assemblies. Bioinformatics 27:2957. 10.1093/bioinformatics/btr507

26. Bolyen E, Rideout JR, Dillon MR, et al (2019) Reproducible, interactive, scalable and extensible microbiome data science using QIIME 2. Nat Biotechnol 37:852–857. 10.1038/s41587-019-0209-9

27. Callahan BJ, McMurdie PJ, Rosen MJ, et al (2016) DADA2: High-resolution sample inference from Illumina amplicon data. Nat Methods 13:581–583. 10.1038/nmeth.3869

28. Conte J, Potoczniak MJ, Tobe SS (2018) Using synthetic oligonucleotides as standards in probe-based qPCR. Biotechniques 64:177–179. doi: 10.2144/btn-2018-2000.

29. Pruesse E, Quast C, Knittel K, et al (2007) SILVA: a comprehensive online resource for quality checked and aligned ribosomal RNA sequence data compatible with ARB. Nucleic Acids Res 35:7188–7196. 10.1093/nar/gkm864

30. Daisley BA, Reid G (2021) BEExact: a Metataxonomic database tool for high-resolution inference of bee-associated microbial communities. mSystems 6:e00082–21. 10.1128/mSystems.00082-21

31. Sf A, Tl M, Aa S, et al (1997) Gapped BLAST and PSI-BLAST: a new generation of protein database search programs. Nucleic acids Res 25(17):3389–402. 10.1093/nar/25.17.3389

32. Johnson M, Zaretskaya I, Raytselis Y, et al (2008) NCBI BLAST: a better web interface. Nucleic Acids Res 36:W5–W9. 10.1093/nar/gkn201

33. Lozupone C, Knight R (2005) UniFrac: a new phylogenetic method for comparing microbial communities. Appl Environ Microbiol 71:8228–8235. 10.1128/AEM.71.12.8228-8235.2005

34. Lozupone C, Lladser ME, Knights D, et al (2011) UniFrac: an effective distance metric for microbial community comparison. ISME J 5:169–172. 10.1038/ismej.2010.133

35. Bisanz JE (2018) qiime2R: Importing QIIME2 artifacts and associated data into R sessions. Version 099 13: https://github.com/jbisanz/qiime2R

36. Żebracka A, Chmielowiec-Korzeniowska A, Nowakowicz-Dębek B, et al (2022) Intestinal microbiota of honey bees (Apis mellifera) treated with amitraz. J Apicult Sci 66:199–207. 10.2478/jas-2022-0015

37. Olofsson TC, Vásquez A (2008) Detection and identification of a novel lactic acid bacterial flora within the honey stomach of the honeybee Apis mellifera. Curr Microbiol 57:356–363. 10.1007/s00284-008-9202-0

38. Olofsson TC, Alsterfjord M, Nilson B, et al (2014) Lactobacillus apinorum sp. nov., Lactobacillus mellifer sp. nov., Lactobacillus mellis sp. nov., Lactobacillus melliventris sp. nov., Lactobacillus kimbladii sp. nov., Lactobacillus helsingborgensis sp. nov. and Lactobacillus kullabergensis sp. nov., isolated from the honey stomach of the honeybee Apis mellifera. Int J Syst Evol Microbiol 64:3109. 10.1099/ijs.0.059600-0

39. Forsgren E, Olofsson TC, Váasquez A, Fries I (2010) Novel lactic acid bacteria inhibiting Paenibacillus larvae in honey bee larvae. Apidologie 41:99–108. 10.1051/apido/2009065

40. Vásquez A, Forsgren E, Fries I, et al (2012) Symbionts as major modulators of insect health: lactic acid bacteria and honeybees. PLoS ONE 7:e33188. 10.1371/journal.pone.0033188

41. Stephan JG, Lamei S, Pettis JS, et al (2019) Honeybee-specific lactic acid bacterium supplements have no effect on American Foulbrood-infected honeybee colonies. Appl Environ Microbiol 85:e00606–19. 10.1128/AEM.00606-19

42. Daisley BA, Pitek AP, Chmiel JA, et al (2020) Novel probiotic approach to counter Paenibacillus larvae infection in honey bees. ISME J 14:476. 10.1038/s41396-019-0541-6

43. Daisley BA, Pitek AP, Chmiel JA, et al (2020) Lactobacillus spp. attenuate antibiotic-induced immune and microbiota dysregulation in honey bees. Commun Biol 3:534. 10.1038/s42003-020-01259-8

44. Daisley BA, Pitek AP, Torres C, et al (2023) Delivery mechanism can enhance probiotic activity against honey bee pathogens. ISME J 17:1382–1395. 10.1038/s41396-023-01422-z

45. Raymann K (2021) Honey bee microbiota and the physiology of antimicrobial resistance. Honey bee medicine for the veterinary practitioner. John Wiley & Sons, Ltd, pp 125–134

46. Bodie A, Corby-Edwards A, Cornell A (2021) Regulation of Dietary Supplements: Background and Issues for Congress. Congressional Research Service. https://sgp.fas.org/crs/misc/R43062.pdf

47. Damico ME (2022) Regulating probiotic use and improving veterinary care to bolster honeybee health. Day One Project. https://www.dayoneproject.org/ideas/regulating-probiotic-use-and-improving-veterinary-care-to-bolster-honeybee-health/

