## Supplemental Figures for "Testing the effectiveness of a commercially sold probiotic on restoring the gut microbiota of honey bees: a field study"

### Supplementary material

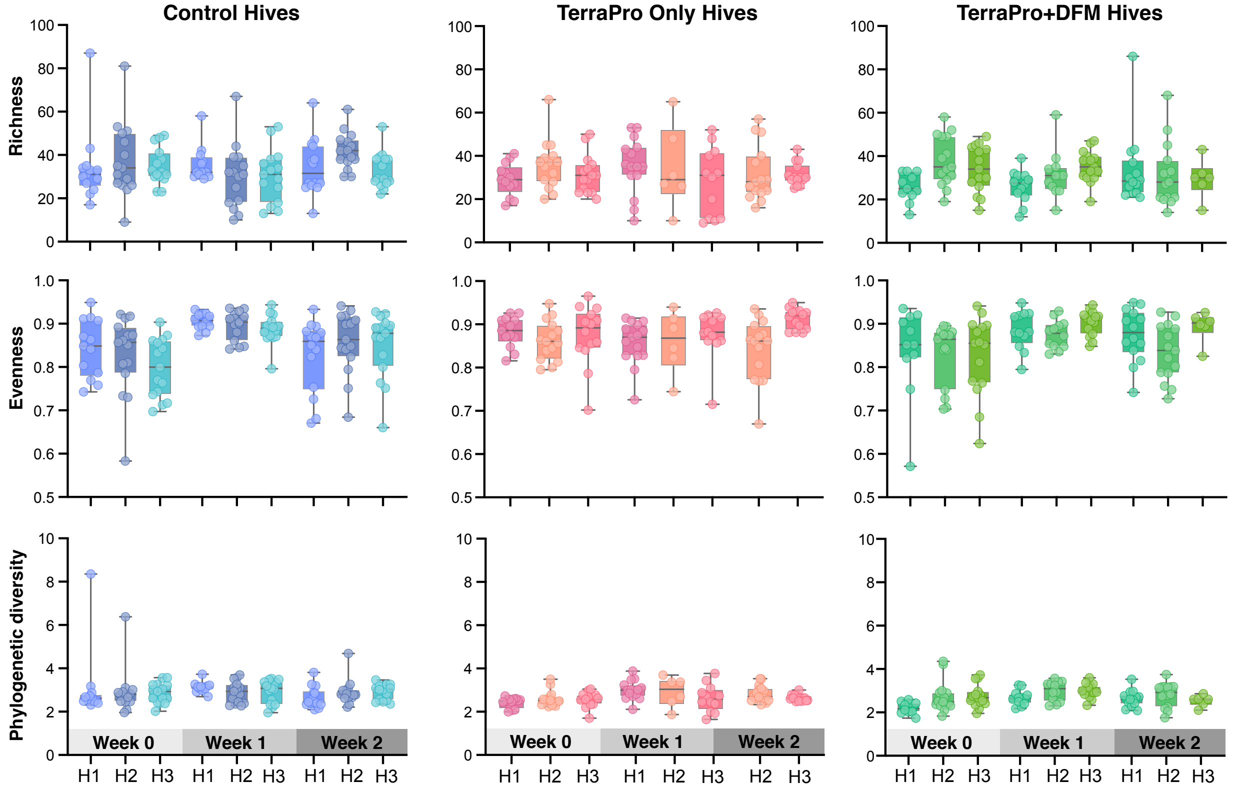


**Figure S1.** Alpha diversity comparisons of the gut microbiota of Control, Terra-Pro-Only, and Terra-Pro+DFM group hives at Week-0 post-Terra-Pro pre-DFM, Week-1 post-Terra-Pro post-DFM, and Week-2 post-Terra-Pro post-DFM. Alpha diversity metrics were based on richness (# of ASVs), evenness (Pielou's Evenness index), and phylogenetic diversity (Faith's Phylogenetic Diversity index). Alpha diversity significance (*P-value*) was determined using the Kruskall-Wallis test with Benjamini-Hochberg FDR correction (*Q-value*), see Dataset S1 for *P* and *Q* values.

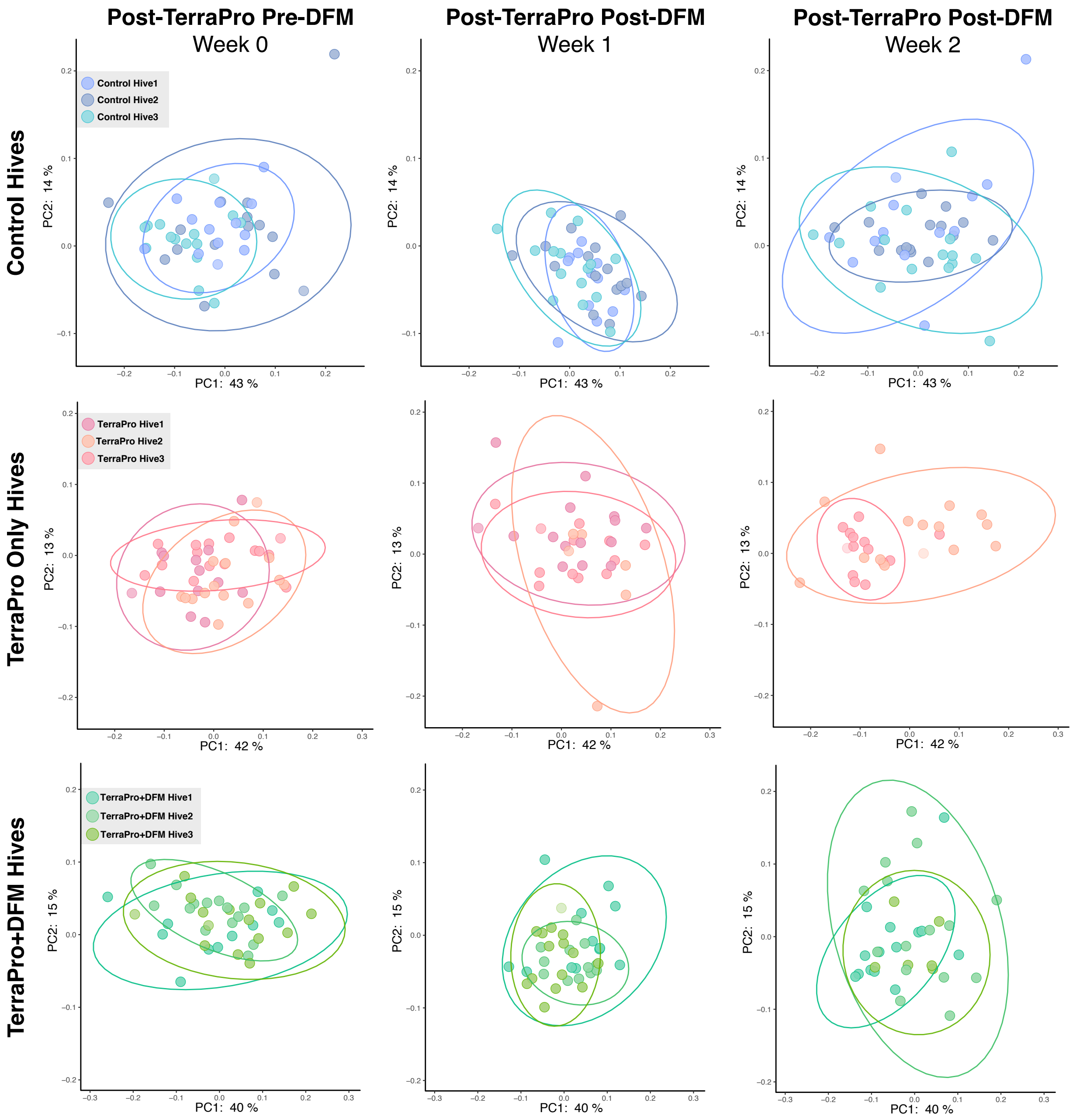


**Figure S2.** Beta diversity comparisons of the gut microbiota of control, Terra-Pro-Only, and Terra-Pro+DFM bees at Week-0 post-Terra-Pro pre-DFM, Week-1 post-Terra-Pro post-DFM, and Week-2 post-Terra-Pro post-DFM. Beta diversity was based on community similarity (weighted UniFrac) and visualized via principal coordinate analysis (PCoA) plots. Significance was tested using PERMANOVA with 999 permutations followed by Benjamini-Hochberg FDR correction (see Dataset S1 for *P* and *Q* values). Ellipses represent the 95% confidence interval.

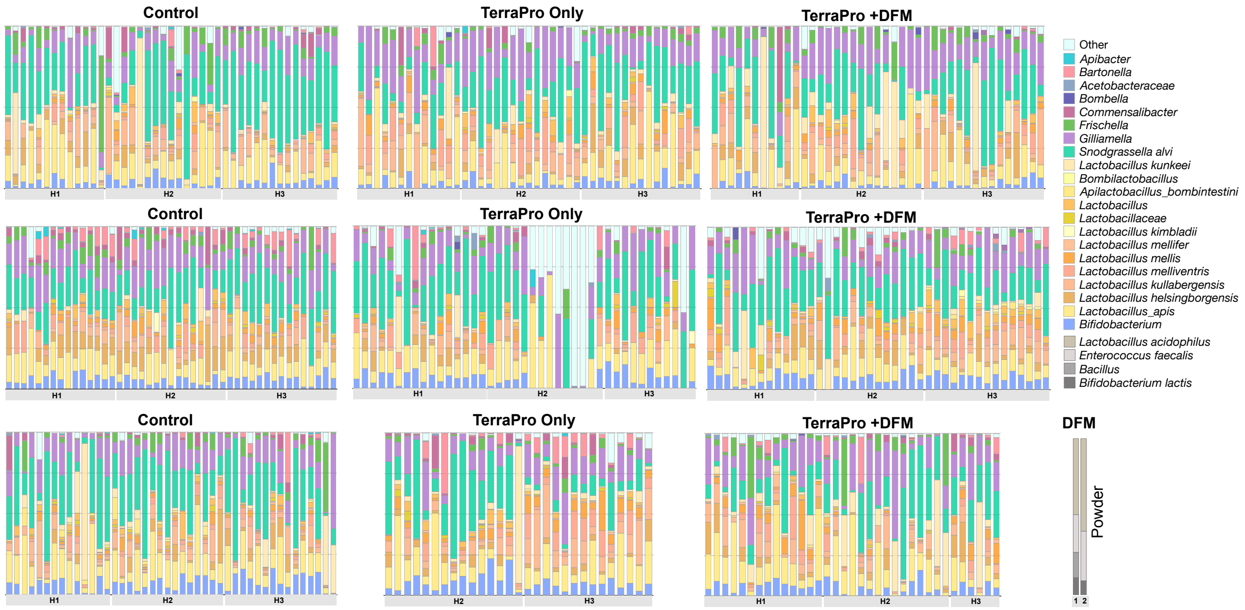


**Figure S3.** Bar plots representing the relative abundance of bacterial taxa found in individual bee samples from each experimental group, hive, and sampling timepoint. The DFM powder (probiotic) was also sequenced as a positive control.
